# Intraspecific diversification and mitonuclear discordance in native versus introduced areas: co-introduction of *Dolicirroplectanum lacustre*, a monogenean gill parasite of the invasive Nile perch *Lates niloticus*

**DOI:** 10.1101/2022.02.26.482128

**Authors:** Kelly J. M. Thys, Maarten P.M. Vanhove, Jonas W. J. Custers, Nathan Vranken, Maarten Van Steenberge, Nikol Kmentová

## Abstract

The Nile perch (*Lates niloticus*) is a notorious invasive species. The introductions of Nile perch into several lakes and rivers in the Lake Victoria region led to the impoverishment of the trophic food webs, particularly well documented in Lake Victoria. Along with the introductions of the Nile perch, its parasites were co-introduced. *Dolicirroplectanum lacustre* (Monogenea, Diplectanidae) is a gill parasite of latid fishes (*Lates* spp.) inhabiting several major African freshwater systems. We examined the intra-specific diversification of *D. lacustre* from *L. niloticus* in Lake Albert (native range) and Lake Victoria (introduced range) by assessing morphological and genetic differentiation, and microhabitat preference. We expected reduced morphological and genetic diversity for *D. lacustre* in Lake Victoria compared to Lake Albert, as a result of the historical introductions. *Dolicirroplectanum lacustre* displays high morphological variability within and between African freshwaters. Mitonuclear discordance within the morphotypes of *D. lacustre* indicates an incomplete reproductive barrier between the morphotypes. The diversification in the mitochondrial gene portion is directly linked with the morphotypes, while the nuclear gene portions indicate conspecificity. Based on our results, we reported reduced genetic and morphological diversity, potentially being a result of a founder effect in Lake Victoria.

## 1. Introduction

The Nile perch *Lates niloticus* (Linnaeus 1758) is considered one of the world’s most invasive species by the Invasive Species Specialist Group of the International Union for Conservation of Nature (IUCN) (Lowe et al. 2000). The introductions of the non-native Nile perch into several of the lakes and rivers in the Lake Victoria region, led to the reduction of species and functional diversity, and as a result dramatically restructured the ecology of the area (Pringle, 2011). These introductions were particularly well documented in Lake Victoria (Ogutu-Ohwayo and Hecky, 1991; Pringle, 2005; Reynolds et al., 1995). In the 1950s and 1960s, the Nile perch was repeatedly introduced into Lake Victoria by the release of captured individuals from Lake Albert and Lake Turkana (Pringle, 2005). In addition to the introductions of non-native tilapias, eutrophication, and overfishing, the introductions of predatory Nile perch lead to the depletion of most, and the extinction of some endemic fish species, causing impoverishment of the trophic food webs (Goudswaard et al., 2006; Ogutu-Ohwayo and Hecky, 1991; Seehausen et al., 1997; Van Zwieten et al., 2016). The introductions transformed the local artisanal fishery in Lake Victoria into a global industrial fishery, with dramatic consequences for the local livelihoods (Pringle, 2011).

Following Pringle (2005), the source population of Nile perch is located in Lake Albert. In Lake Albert, *L. niloticus* occurs in sympatry with *Lates macrophthalmus* Worthington, 1929. While *L. niloticus* is larger and mainly occurs inshore, *L. macrophthalmus* is a smaller, offshore species (Holden, 1967). The limited genetic evidence based on allozymes suggests that these are indeed distinct species, and that Nile perch in Lake Victoria are largely represented by individuals of *L. niloticus* from Lake Albert, but the presence of *L. macrophthalmus* in Lake Victoria cannot be excluded (Hauser et al. 1998).

The threats of alien species have been highlighted many times, but the co-introduction of their possibly co-invasive parasites deserves more attention, as its consequences are usually underestimated (Kmentová et al., 2018; Lymbery et al., 2014; Peeler et al., 2004; Šimková et al., 2018). *Co-introduced* parasites are parasites that have been transported with an alien host to a new locality outside their natural range. There, they become established by survival, reproduction, and dispersal within the alien hosts. In case parasites were transported but have not established in the introduced range, it is unlikely they are ever recorded (Lymbery et al. 2014). *Co-invasive* parasites are those that have been co-introduced, and then have spread to new, native hosts (Lymbery et al., 2014), so-called spill-over events (Prenter et al. 2004; Goedknegt et al. 2016, 2017). The invasive potential, and success of parasite establishment is affected by several ecological factors, such as the size of the founder population of the host and consequently that of the parasite (Anderson and May, 1991; Dlugosch and Parker, 2008; Sakai et al., 2001), the parasite’s life cycle, and environmental biotic and abiotic conditions (Lymbery et al., 2014; Taraschewski, 2006). Hence, co-invasion is not a straightforward consequence of co-introduction. Along with the introductions of the Nile perch, some of its parasites were co-introduced, more specifically, the monogenean (Platyhelminthes, Monogenea) parasite *Dolicirroplectanum lacustre* (Thurston & Paperna 1969), and possibly, the copepods *Ergasilus kandti* Van Douwe 1912 and *Dolops ranarum* (Stuhlmann 1892) (Thurston and Paperna 1969), the myxosporean *Henneguya ghaffari* Ali 1999, and the nematodes *Contracaecum multipapillatum* (Drasche 1882) and *Cucullanus* spp. Müller, 1777 (Outa et al., 2021).

Monogeneans are parasitic flatworms that can be identified morphologically by their hard parts: the attachment organ (haptor) and the male copulatory organ (MCO) (Woo, 2006). These ectoparasites have a direct life cycle in which short-living, ciliated larvae (oncomiracidia) hatch from eggs and attach to their single host, presumably synchronised with host behaviour (Kearn, 1973). More specifically, members of Diplectanidae metamorphose on their single host into an adult with a functional male reproductive system, and later a fully mature hermaphrodite that delivers eggs after cross-fertilisation (Whittington et al. 1999). Monogeneans usually exhibit high host specificity (i.e., restricted to one host group or host species) (Odening, 1976). Exceptions have been recorded, such as *Neobenedenia melleni* (MacCallum, 1927), a pathogenic species which has the broadest host specificity of all monogenean species, recorded from over 100 host species (Bullard et al., 2000; Whittington and Horton, 1996).

Various speciation processes (e.g., co-speciation and intrahost speciation) are proposed to affect monogenean diversity (Šimková et al., 2004; Vanhove et al., 2015). Parasites tend to diversify faster than their hosts due to their short generation times and faster mutation rates (Nieberding et al., 2004; Nieberding and Olivieri, 2007). This should be especially apparent in comparison with long-lived and large hosts, as would be expected for *D. lacustre* from its latid hosts (Kmentová et al., 2020a). The study of parasite population structure may increase the resolution for understanding host population structure, phylogeography (Nieberding et al., 2004), and introduction pathways (Ondračková et al., 2012). Following (Buchmann and Lindenstrøm, 2002), host species selection is a consequence of dynamic interactions between the parasite and its host over time. Selection acts on the scale of microhabitats on the gills of a single host species, through differential microhabitat preferences (Koskivaara et al. 1992; Buchmann and Uldal 1997; Raymond et al. 2006). Purportedly, the morphology of the attachment organs is an important adaptation to the monogenean’s host, and to specific microhabitats within the host, since these structures could influence the ability of the parasite to infect, feed, and reproduce on a specific host species (Šimková et al., 2002).

*Dolicirroplectanum lacustre* (Monogenea, Diplectanidae) is the single known monogenean gill parasite of four species of lates perches (Perciformes, Latidae) in African freshwaters (*L. niloticus, Lates microlepis* Boulenger 1898*, Lates angustifrons* Boulenger 1906, and *Lates mariae* Steindachner 1909) that have been examined for parasites (Kmentová et al., 2020a). *Dolicirroplectanum lacustre* displays a high morphological variation: a “wide range of shapes and sizes” (Thurston and Paperna, 1969). In the original description of *D. lacustre*, Thurston and Paperna (1969) noted these differences and identified a so-called ‘slender form’, with a well delineated haptor at the posterior end of the body, and a usually longer ‘gravid form’ which is proportionally wider and has an embedded haptor. In accordance with Thurston and Paperna (1969), Kmentová et al. (2020a) reported distinct morphotypes of *D. lacustre* in Lake Albert and continuous morphological variation across several African freshwater systems across a broad geographic scale, indicating phenotypic plasticity of the species. Although *D. lacustre* showed high morphological variation, the genetic differentiation between parasites of allopatric host species was not as high as typically associated with distinct diplectanid species (Kmentová et al., 2020a).

In this study, we investigate the fine-scale morphological and genetic differentiation of *D. lacustre* from Nile perch in its native (Lake Albert) and introduced range (Lake Victoria). We hypothesise that: (1) a founder effect had taken place in Lake Victoria. Hence, we expect reduced genetic and morphological diversity within *D. lacustre* in its introduced range in comparison with its native population in Lake Albert; (2) we expect high phenotypic variation and low genetic differentiation in Lake Albert, as supported by the presence of distinct morphotypes in earlier studies (Kmentová et al., 2020a; Thurston and Paperna, 1969); (3) there has been a niche shift among the populations in Lake Albert that resulted in differential gill microhabitat preferences which led to morphological changes of *D. lacustre,* producing an (imperfect) reproductive barrier between the morphotypes.

## 2. Materials and methods

### 2.1 Sampling of the Nile perch and its gill parasites

Fish samples of *L. niloticus* from Lake Victoria and Lake Albert were examined (Table 1). Fresh specimens were obtained from local fishermen in Kaiso, Uganda (Lake Albert) and Kikondo, Uganda (Lake Victoria) during a field expedition in 2019 (Figure 1). In total, the gills of 15 fish specimens (8 samples from Lake Albert, 7 samples from Lake Victoria) were dissected and preserved in ethanol (99% EtOH). The pairs of gills from 12 randomly chosen specimens (5 specimens from Lake Albert, 7 specimens from Lake Victoria) were examined following the standard protocol of Ergens and Lom (1970) using a Leica EZ4 stereomicroscope. Monogenean gill parasites were extracted, some of the individuals were cut in three parts with the posterior and anterior parts mounted on slides (using Hoyer’s medium as fixative) for morphological characterisation, and the central part preserved in ethanol (99% EtOH) for genetic analyses. In addition, several individuals were mounted on slides as entire specimens and fixed (Hoyer’s medium). Whole mounted parasite specimens were drawn by group according to locality and morphotype (Thurston & Paperna 1969): Lake Albert (gravid), Lake Albert (slender), Lake Victoria using a Leica DM2500 optical microscope at a magnification of 1000x (objective x100 1.32 oil XHC PL FLUOTAR, ocular x10) and a reMarkable^®^ graphical tablet. Drawings were edited in Adobe Photoshop® 2021 (v 22.3.1).

**Fig. 1.**
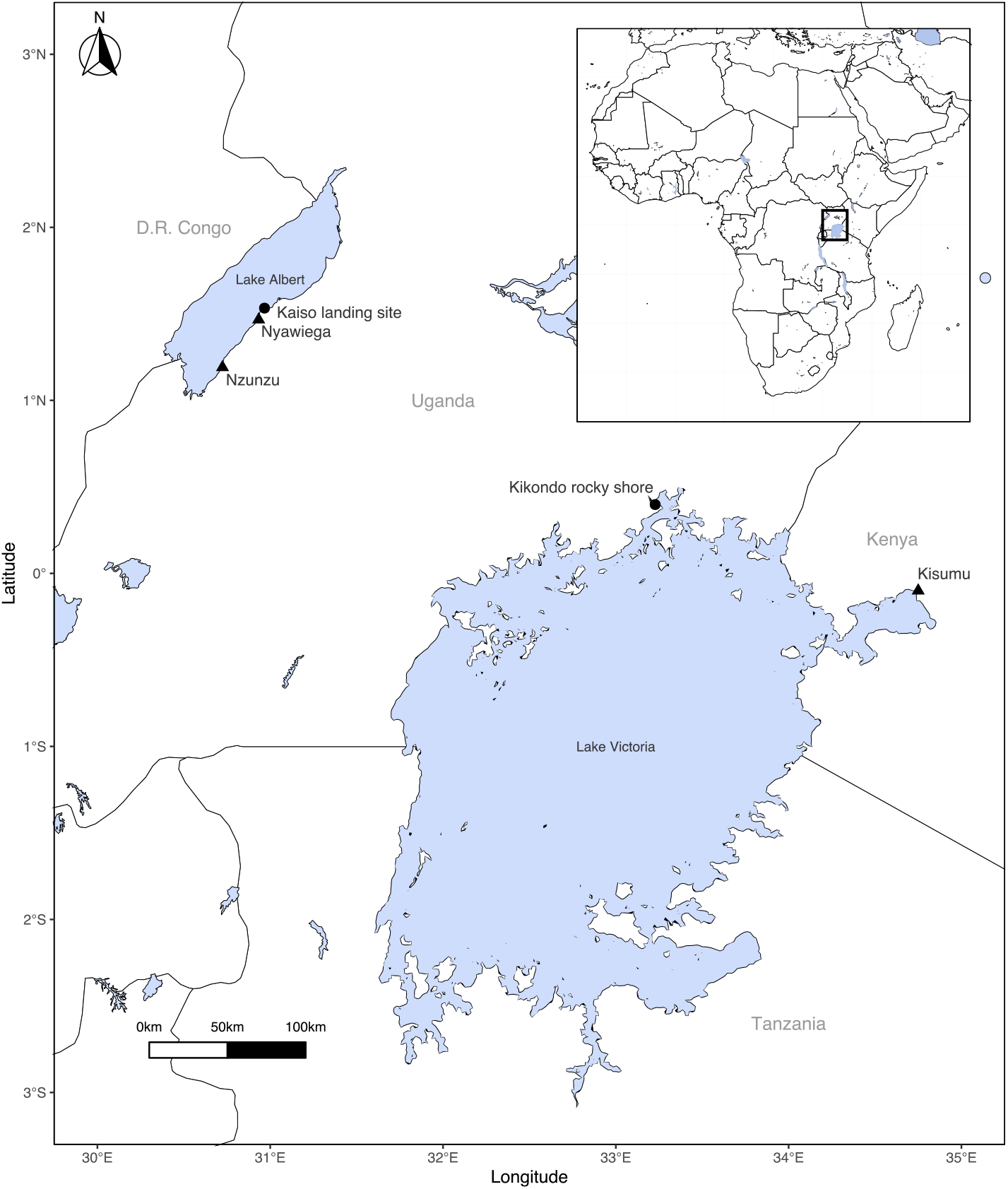
Sampling localities of *Lates niloticus* at Lake Albert and Lake Victoria: Kaiso landing site, Lake Albert, Uganda (N 00°23’52.0’’, E 33°13’39.6”) (n = 8) and Kikondo rocky shore, Lake Victoria, Uganda (N 01°31’59.3’’, E 30°58’00.9”) (n = 7). Sampled localities from Kmentová et al. (2020a) depicted by triangles: Kisumu, Lake Victoria, Kenya (00°06’S, 34°45’E; n = 3) and Nzunzu, Lake Albert, Uganda (N 01°11’27.6’’, E 30°43’26.4”; n = 12) and Nyawiega, Lake Albert (01°28’N, 30°56’E; n = 3). We used the geographical databases ‘Global Lakes and Wetlands Database’ (WWF, Lehner and Dooll) and the ‘High Resolution World Vector Map’ from package rnaturalearth (South 2021) to map the lakes, and countries respectively. The map was created using the packages sf (Pebesma 2018), ggspatial (Dunnington 2021) and ggplot 2 (Wickham 2016) in R Studio v 1.4.1106 (RStudio Team 2020).

**Table 1.**
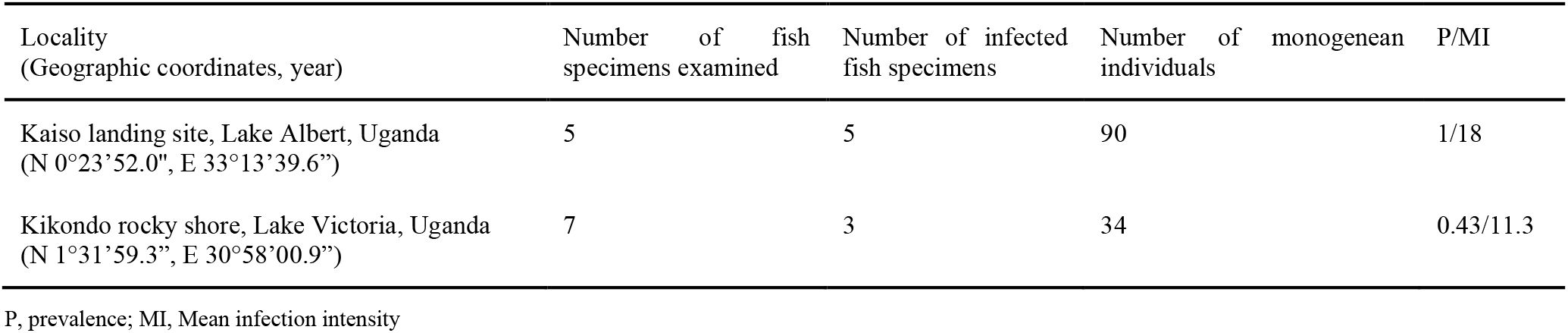
Overview of host species examined for monogenean parasites and infection parameters per locality.

Infection parameters prevalence (P) and mean infection intensity (MI) were calculated following Bush et al. (1997). Prevalence was determined as the proportion of host specimens infected with *D. lacustre*. The mean infection intensity was calculated as the ratio of the total number of parasite specimens of *D. lacustre* to the total number of host specimens infected by *D. lacustre*. During extraction of the parasites, the microhabitat of each parasite specimen on the gill was recorded by subdivision of each gill arch into nine microhabitats. Following Gobbin et al. (2021) each gill arch was subdivided along the longitudinal axis (dorsal, median, ventral) and transversal axis (proximal, central, distal; from gill bar to tips of gill filaments).

### 2.2 Morphometrics of the parasites

Measurements of the hard parts, total body size and distances between the two pairs of eyespots were obtained from 71 specimens at a magnification of 1000x (objective x100/1.32 oil XHC PL FLUOTAR, ocular x10) with differential interference contrast on a Leica DM2500 optical microscope using Las X software v3.6.0.20104. A total of 20 parameters of the hard parts of the haptor and MCO were measured (Figure 2). The terminology of (Justine and Henry, 2010) was followed. To investigate morphological differentiation in haptor morphology between localities and morphotypes, raw haptor measurements were analysed by multivariate statistical techniques in RStudio® (R Core Team 2020). A Principal Component Analysis (PCA) was performed with 23 measurements from 56 individuals (combined with measurements of 22 individuals from Kmentová et al. (2020)) using the package FactoMineR (Lê et al. 2008). Individuals were selected for the PCA when more than 50% of the measured characteristics were obtained. Missing values for measurements were imputed by the variable mean as default in the package factoextra (Kassambara and Mundt 2017). Results were visualised using the packages ggplot2 (Wickham, 2016), reshape2 (Wickham, 2007), factoextra (Kassambara and Mundt 2017), tidyr (Wickham et al. 2021), and MASS (Venables & Ripley 2002).

**Fig. 2.**
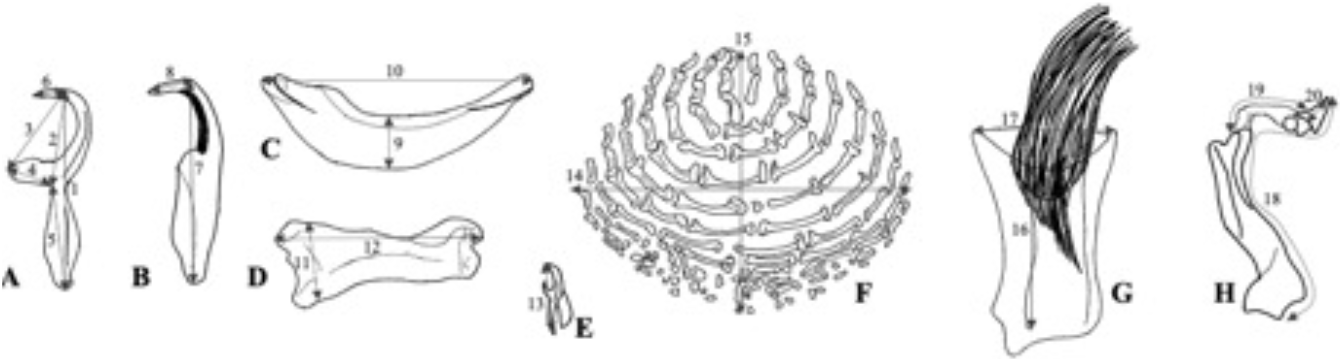
Measurements for sclerotized structures of haptor and reproductive organs of *Dolicirroplectanum lacustre*: A, Ventral anchor: 1, total length; 2, length to notch; 3, length to inner root, 4, inner root length, 5, outer root length, 6, point length; B - Dorsal anchor: 7, total length, 8, point length; C - Ventral bar: 9, maximum width, 10, straight length; D - Dorsal bar: 11, maximum width, 12, straight length; E - Hook: 13, hook length; F - Squamodisc: 14, squamodisc width; 15, squamodisc length; G - Male copulatory organ: 16, copulatory tube length; 17, copulatory tube width; H - Vagina: 18, total length; 19, tube length; 20, point length. Structures drawn of *D. lacustre* from host specimen HP4318 from Lake Victoria.

To investigate the relationship between groups per lake and morphotype, and level of differentiation in the measurements of the MCO, parametric tests were applied. Analysis of Variance (ANOVA) was applied for overall effects; for pairwise comparisons, unpaired t-tests were applied, implemented in the package stats (R Core Team 2020). Assumptions of normality and homogeneity of variance were tested by Shapiro-Wilk’s W tests from the package stats (R Core Team 2020) and Levene’s tests from the car package (Fox and Weisberg 2019), respectively, as well as by graphical examination of the residuals.

### 2.3 Molecular characterisation of the parasites

Whole genomic DNA was extracted from the central part of 64 monogenean parasite individuals (20 from Lake Victoria, 44 from Lake Albert) that were preserved in 99% ethanol using the following protocol. The sample was spinned down and ethanol was removed. 195 μL of TNES buffer (400 mM NaCl, 20 mM EDTA, 50 mM Tris (pH 8, 0.5% SDS)), and 5 μL of proteinase *K* (20 mg/mL) were added to the sample. After 45 min of incubation at 55 °C, 2 μL of Invitrogen™ yeast RNA (10 mg/mL) was added as a carrier. Next, 65 μL of NaCl (5 M) and 290 μL of 96% ethanol were added, and the sample was cooled for 60 min at −20 °C. The sample was spinned down for 15 min at 18,000 rcf to a small pellet. Supernatant was removed and substituted by 1 mL of chilled 70% ethanol. Next, the samples were centrifuged for 5 min at 18,000 rcf. The ethanol rinse step (removal of supernatant, addition of ethanol and centrifugation) was repeated once again. The ethanol was removed, and the DNA was eluted in 100 μL of 0.1X TE (0.02 % tween-20). The DNA extract was placed at 4 °C for resuspension overnight, and stored at a temperature of −20°C.

Sequences of a portion of the mitochondrial cytochrome *c* oxidase subunit 1 (COI) gene, and nuclear gene portions from the small and large ribosomal subunit gene (18S rDNA and 28S rDNA) and internal transcriber spacer 1 (ITS-1) were obtained following PCR. The three nuclear markers evolve at different rates and are suitable to assess genetic divergence at the interspecific level (Vanhove et al., 2013). COI is the current standard marker for intraspecific genetic differentiation in many monogenean taxa (Kmentová et al., 2020b) due to its relatively high rate of molecular evolution in comparison to rDNA. Part of the mitochondrial COI gene was amplified using ASmit1 (5’-TTTTTTGGGCATCCTGAGGTTTAT-3’) (Littlewood et al., 1997) combined with Schisto3 (5’-TAATGCATMGGAAAAAAACA-3’) (Lockyer et al., 2003), and with Asmit2 (5’-TAAAGAAAGAACATAATGAAAATG-3’) (Littlewood et al., 1997) in a nested PCR following Vanhove et al. (2015). For both primer combinations, the amplification reaction contained 24 μL of PCR mix (0.2 μL of 1 unit Invitrogen Taq polymerase, 2.5 μL Invitrogen PCR buffer, 2 μL of 2 mM MgCl2, 0.5 μL of 0.2 mM dNTPs, 2μL of 0.8 μM of each primer, 14.80 μL of dd H2O) with 1 μL of isolated DNA (concentration was not measured) in a total reaction volume of 25 μL and was performed under the following conditions for the first reaction: initial denaturation at 95 °C for 5 min, 40 cycles of 1 min at 94 °C, 1 min at 44 °C and 1 min at 72 °C, and a final 7 min at 72 °C. The second (nested) reaction was performed under the following conditions: 5 min at 95 °C, 40 cycles of 1 min at 94 °C, 1 min at 50 °C and 1 min at 72 °C, and a final 7 min at 72 °C. Primers C1 (5’-ACCCGCTGAATTTAAGCAT-3’) and D2 (5’-TGGTCCGTGTTTCAAGAC-3’) (Hassouna et al., 1984) were used for amplification of the partial 28S rDNA gene. Each PCR reaction contained 1 unit of Q5® Hot Start High-Fidelity DNA Polymerase, 1X PCR buffer containing 0.1 mg/mL bovine serum albumin (BSA), 0.2 mM dNTPs, 0.5 μM of each primer and 2 μL of isolated DNA (concentration not measured) in a total reaction volume of 25 μL. The reaction proceeded under the following conditions: 2 min at 95 °C, 39 cycles of 20 s at 94 °C, 30 s at 65 °C, 1 min 30 s at 72 °C, and 10 min. at 72 °C. Partial 18S rDNA and ITS-1 were amplified using the S1 (5’-ATTCCGATAACGAACGAGACT-3’) (Sinnappah et al., 2001) and IR8 (5’-GCTAGCTGCGTTCTTCATCGA-3’) (Šimková et al., 2003) primers. Each reaction contained 1 unit of Q5® Hot Start High-Fidelity DNA Polymerase, 1X PCR buffer containing 0.1 mg/mL BSA, 0.2 mM dNTPs, 0.5 μM of each primer and 2 μL of isolated DNA (concentration not measured) in a total reaction volume of 25 μL under the following conditions: 2 min at 95 °C, 39 cycles of 1 min at 94 °C, 1 min at 64 °C and 1 min 30 s at 72 °C, and a final elongation of 10 min at 72 °C. PCR amplification success was verified by agarose gel electrophoresis. A volume of 10 μl of the PCR product was enzymatically purified for positive samples, using 4 μL of ExoSAP-IT reagent under the following conditions: 15 min at a temperature of 37 °C, and 15 min at 80 °C. Purified PCR products were sent out to Macrogen Europe B.V. for Sanger sequencing; amplification primers were used for sequencing reactions. The acquired sequences for each marker were visually inspected and trimmed using the software Geneious v2021.1.1. Sequences were aligned with previously published sequences of *D. lacustre* from Kmentová et al. (2020a) using MUSCLE (Edgar, 2004) under default settings as implemented in Geneious v2021.1.1.

### 2.4 Genetic differentiation and phylogeography

Per marker, uncorrected pairwise genetic distances (p-distances) among sequences were computed in Geneious Prime v2021.1.1. Already available sequences of the four genetic markers of *D. lacustre* from Lake Albert (GenBank accession numbers: MK908145.1-MK908196.1, MK937576.1, MK937579.1-MK937581.1, MK937574.1, MK937575.1) were included in the analyses. Haplotype networks were constructed using Median Joining networks following Bandelt et al. (1999) in PopART v1.7104 (Leigh and Bryant, 2015) for each marker separately. The analyses of population structure and demographic history within *D. lacustre* were based on a 325 bp fragment of COI. This allows for the detection of recent evolutionary events, such as possible incipient speciation as a result of host preference (Kmentová et al., 2016). Genetic diversity of COI was assessed by the number of haplotypes and polymorphic sites, haplotype diversity (h) and nucleotide diversity (π), calculated in Arlequin v3.5.2.2 (Excoffier and Lischer, 2010). Differentiation among pre-defined populations (according to locality and morphotype) was estimated by F_ST_. Analysis of Molecular Variance (AMOVA) based on F-statistics was applied to test for a significant population structure of *D. lacustre* at the level of locality (non-native and native). To test for signals of past population expansion, two different neutrality tests, Tajima’s D and Fu’s F_S_ were calculated in Arlequin v3.5.2.2 (Excoffier and Lischer, 2010) for the two hypothetical populations. First, the neutrality tests were calculated for the specimens from Lake Albert. Second, the Lake Victoria individuals were added.

### 2.5 Microhabitat description

Parasite microhabitats were visualised by the combination of the spatial distribution of *D. lacustre* on each pair of gills (left and right gills per host individual) into a single spatial distribution. Per lake/morphotype, the spatial distribution of all individuals was summarised in a heat map using the geom_tile function as part of the ggplot2 package (Wickham, 2016). As replicates per population were limited, no statistical effects could be verified.

## 3. Results

A total of 124 monogenean gill parasites were retrieved from the 8 host specimens examined in this study (Table 1). The parasite specimens were morphologically identified as *Dolicirroplectanum lacustre*. Diplectanid-specific haptoral characteristics included the presence of ventral and dorsal squamodiscs, and three transversal bars connected to pairs of ventral hooks and dorsal hooks. *Dolicirroplectanum lacustre* can be distinguished from other diplectanids by the observed characteristics: a longer outer root in the ventral anchors (Thurston and Paperna, 1969); a wide, robust, and barrel-shaped MCO formed by two anteriorly oriented nested tubes, one encasing the other (Kmentová et al., 2020a); the presence of an accessory piece; squamodiscs composed of variable concentric rows of bone shaped rodlets forming open rings; reduced inner roots of the ventral anchor. Additional characteristics include: a simple prostatic reservoir; seminal vesicle as an expansion of vas deferens; intercaecal, pre-testicular ovary encircles right caecum (Kmentová et al., 2020a; Thurston and Paperna, 1969). The dorsal bars are large, bone-shaped structures with a broad base pointing towards the centre of the haptor. The ventral bar has an elongated, oval shape with tapering ends. A sclerotized vagina was observed in 29 out of the 71 measured specimens. Additional observed characteristics include two pairs of eyespots, of which the posterior eyespots are larger and closer together, a thin tegument covering the body, and 14 rudimentary hooklets of equal sizes.

Infection parameters varied per location, with greater prevalence (P) and mean infection intensity (MI) in Lake Albert compared to Lake Victoria (Table 1). We observed co-infections (infection by both morphotypes of *D. lacustre* on a single host individual of *L. niloticus)* of the morphotypes on two host individuals in Lake Albert (HP4094, HPAlb.X).

### 3.1 Morphotypes and morphometric variation

During the screening procedure, clear size differences in total body length and haptoral sclerotised structures were apparent between individuals of *D. lacustre* from Lake Albert. Therefore, we classified the specimens from Lake Albert under the gravid morphotype (n=9) and slender morphotype (n=42) following Thurston and Paperna (1969). Specimens of the gravid morphotype (Fig. 3a) were identified by the following haptoral features: proportionally larger body, longer ventral anchors with a longer outer root, longer dorsal anchors with a shorter tip, long and narrow ventral and dorsal bars, larger ventral and dorsal squamodiscs (Table 2). The MCO for gravid individuals was longer, as was the vagina (Table 2). Also, the distance between the eyespots of both pairs was larger in the gravid morphotype. Conversely, specimens of the slender morphotype (Fig. 3b) were identified by a proportionally smaller body, shorter ventral anchors with a shorter outer root, shorter dorsal anchors with a longer tip, wider but shorter ventral and dorsal bars, and smaller squamodiscs. The copulatory tube for the slender morphotype was usually shorter, as was the vagina total length and tube length (Table 2; Supplementary data S3). The individuals retrieved from Lake Victoria had sclerotized structures of the slender morphotype (Fig. 3c). However, these individuals were smaller in total body size, they had shorter ventral and dorsal anchors, narrower ventral, and dorsal bars, and squamodiscs were less wide in comparison with both morphotypes in Lake Albert. The specimens from Lake Victoria had a shorter copulatory tube and vagina in comparison with the specimens in Lake Albert (Table 2).

**Fig. 3.**
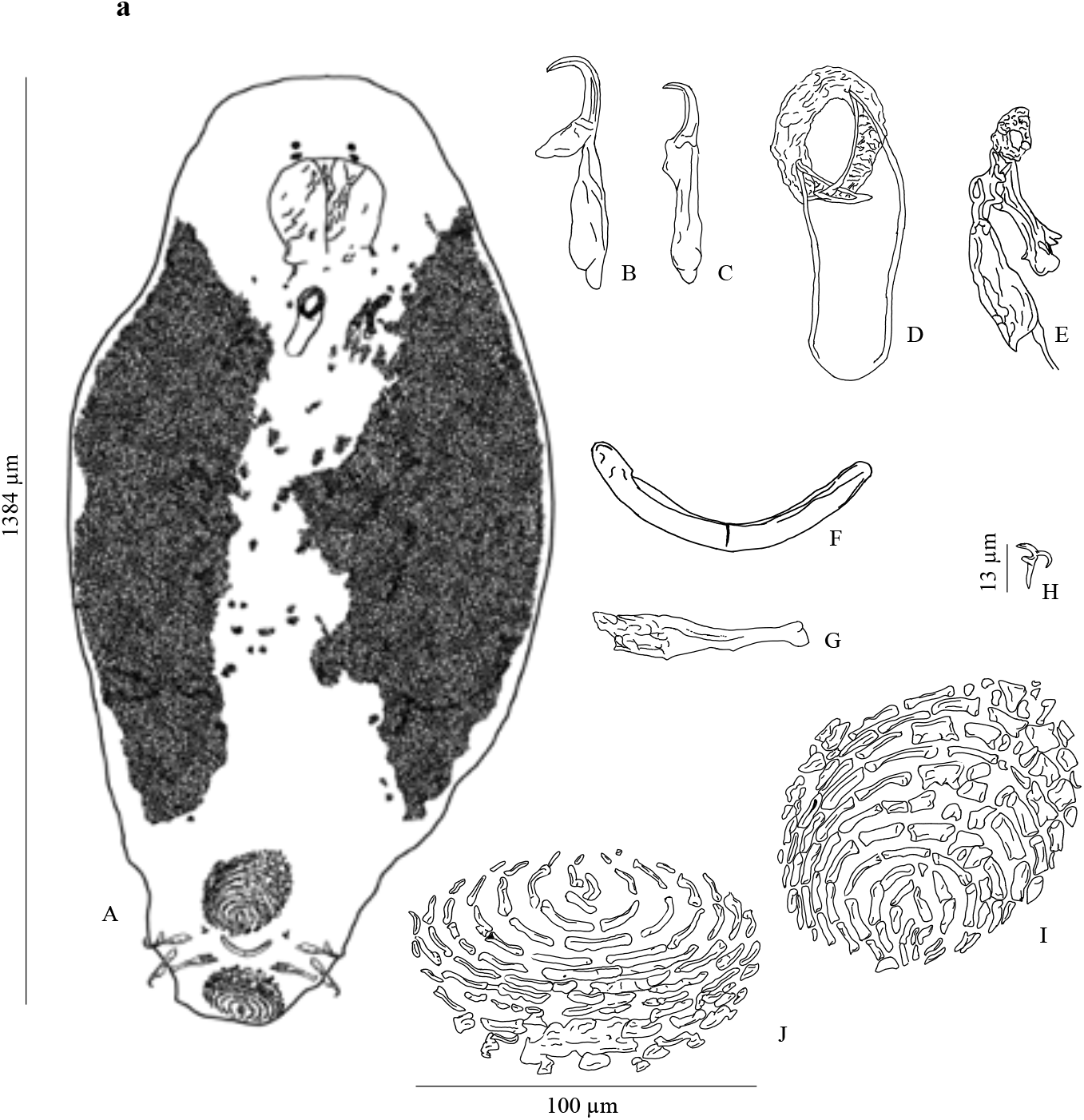
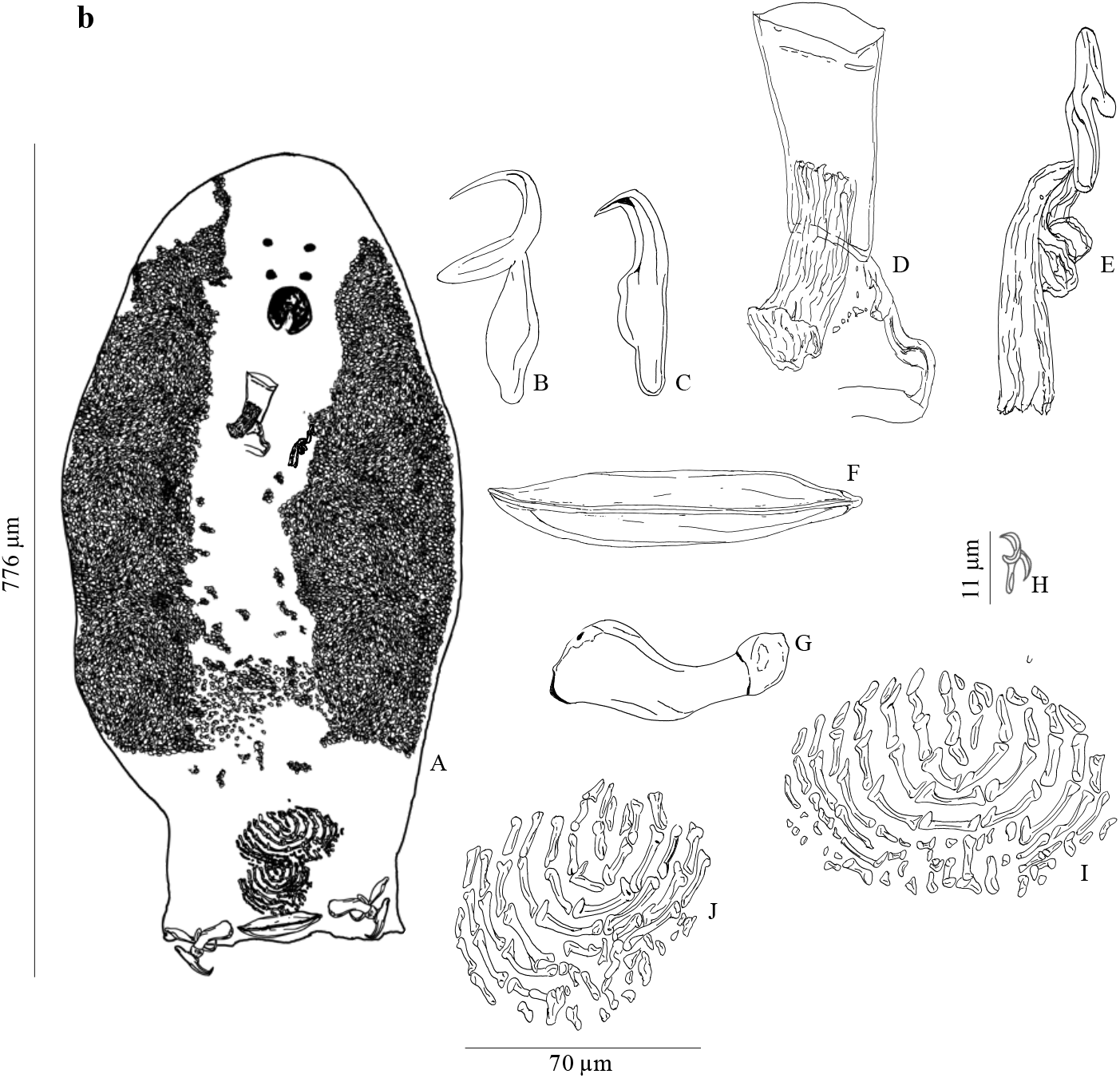
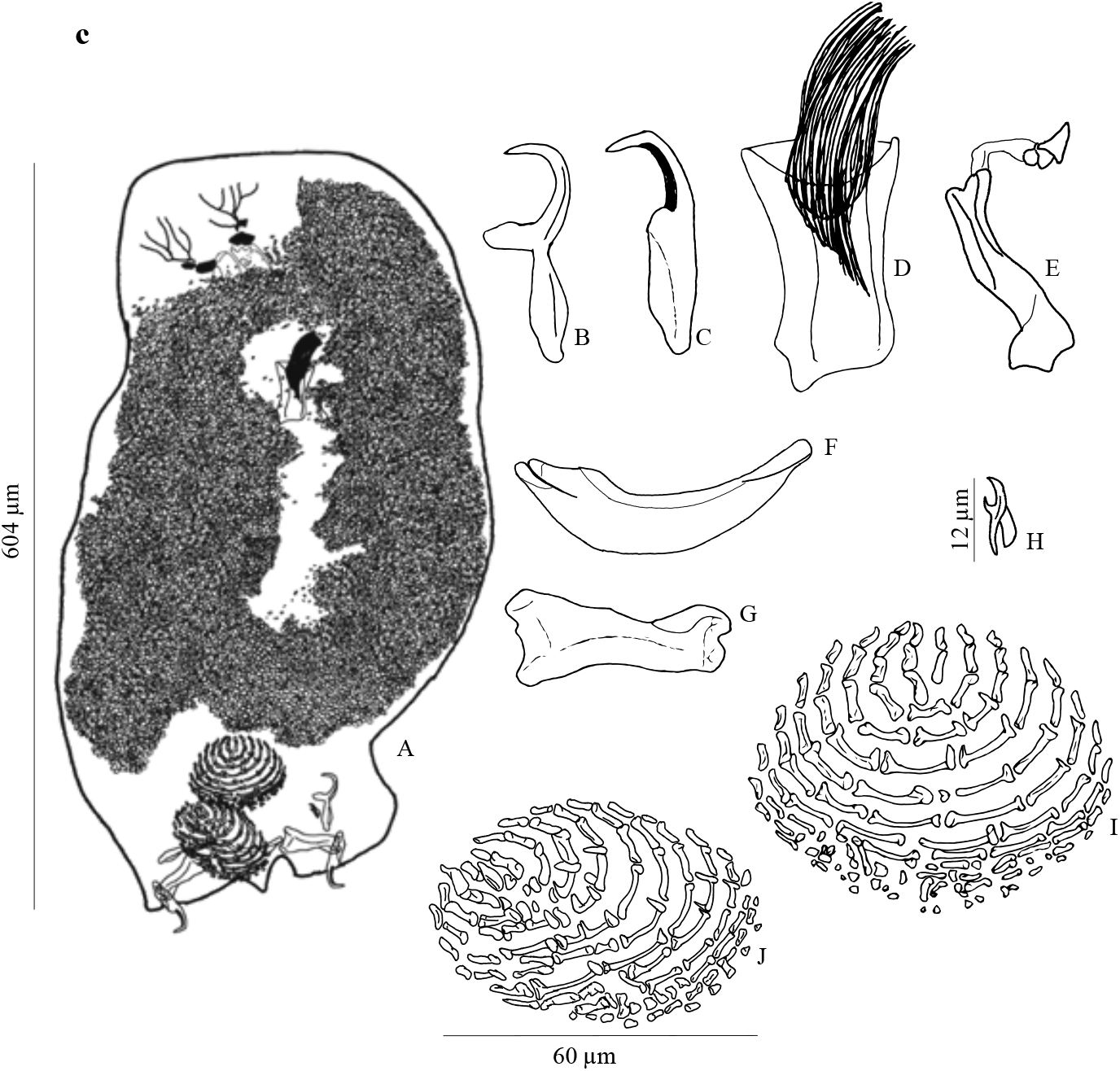
Morphotypes of *Dolicirroplectanum lacustre* collected from *Lates niloticus.* Specimens drawn from a dorsal view with Hoyer’s medium as fixative. a *Dolicirroplectanum lacustre* gravid morphotype from Lake Albert (host specimen HP4094); (b) *Dolicirroplectanum lacustre* slender morphotype from Lake Albert (host specimen HP4099); (c) *Dolicirroplectanum lacustre* from Lake Victoria (host specimen HP4318). Drawings from the dorsal view with A, whole mount; B, ventral anchor; C, dorsal anchor; D, male copulatory organ; E, vagina; F, ventral bar; G, dorsal bar; H, hook; I, ventral squamodisc; J, dorsal squamodisc.

**Table 2.** Overview of morphometric measurements performed on haptoral and genital sclerotised structures of *Dolicirroplectanum lacustre.* Values depict the average measurements ± the standard deviations; min. - max. values, and *n*, the number of individuals.

| Parameters ($\mu\text{m}$) | *L. niloticus*, Lake Victoria | *L. niloticus*, Lake Albert (slender morphotype) | *L. niloticus*, Lake Albert (gravid morphotype) |
| --- | --- | --- | --- |
| Total length | 533.4 ± 47.0 (475.9–603.7); n=6 | 776.6 ± 97.1 (601.3–899.3); n=8 | 1384.0; n=1 |
| Total width | 298.6 ± 61.1 (222.2–417.4); n=12 | 361.6 ± 91.7 (214.6–555.4); n=24 | 947.8 ± 129.5 (843.7–1245.0); n=8 |
| *Ventral anchor* |  |  |  |
| Total length | 47.8 ± 2.4 (44.2–51.6); n=26 | 51.1 ± 3.8 (45.4–59.3); n=25 | 63.2 ± 4.0 (58.9–69.1); n=8 |
| Length to notch | 22.2 ± 0.74 (20.6–23.5); n=26 | 21.1 ± 1.8 (17.9–24.5); n=25 | 24.5 ± 0.8 (23.0–25.7); n=8 |
| Length to inner root | 25.4 ± 1.2 (22.4–27.8); n=26 | 25.5 ± 1.3 (23.1–28.8); n=25 | 28.8 ± 1.3 (26.5–30.4); n=8 |
| Inner root length | 12.4 ± 0.9 (11.2–14.8); n=26 | 12.8 ± 0.7 (11.1–14.0); n=25 | 14.9 ± 0.9 (13.2–16.0); n=8 |
| Outer root length | 28.5 ± 2.2 (22.5–32.9); n=26 | 31.7 ± 3.3 (26.03–37.6); n=25 | 40.9 ± 4.1 (36.5–46.5); n=8 |
| Point length | 9.8 ± 0.9 (8.1–11.9); n=26 | 9.5 ± 0.7 (7.9–11.0); n=25 | 10.6 ± 1.3 (8.5–12.0); n=8 |
| *Dorsal anchor* |  |  |  |
| Total length | 42.3 ± 2.1 (37.2–44.9); n=26 | 45.0 ± 2.6 (39.5–52.6); n=25 | 56.5 ± 3.1 (50.4–59.7); n=8 |
| Point length | 8.6 ± 1.3 (4.0–10.5); n=26 | 9.3 ± 0.9 (7.8–10.5); n=25 | 6.5 ± 0.8 (5.2–8.0); n=8 |
| *Ventral bar* |  |  |  |
| Straight length | 65.7 ± 5.2 (55.7–74.1); n=24 | 68.9 ± 11.7 (37.4–94.6); n=25 | 80.4 ± 10.0 (66.9–95.2); n=8 |
| Maximum width | 16.2 ± 1.5 (11.7–18.1); n=24 | 16.8 ± 2.3 (11.4–19.5); n=25 | 6.1 ± 0.6 (5.5–6.8); n=8 |
| *Dorsal bar* |  |  |  |
| Straight length | 44.8 ± 3.9 (33.1–52.5); n=26 | 51.6 ± 6.1 (34.1–61.5); n=25 | 56.4 ± 6.6 (45.3–61.7); n=8 |
| Maximum width | 16.9 ± 2.1 (13.2–23.0); n=26 | 17.6 ± 2.7 (11.0–22.3); n=25 | 11.4 ± 2.6 (6.7–14.2); n=8 |
| *Ventral squamodisc* |  |  |  |
| Width | 69.3 ± 13.5 (51.0–102.2); n=23 | 80.4 ± 18.9 (50.1–125.4); n=19 | 115.9 ± 29.8 (83.9–156.1); n=8 |
| Length | 63.5 ± 11.9 (43.6–78.9); n=23 | 57.1 ± 14.1 (28.4–86.9); n=19 | 108.7 ± 39.5 (69.5–191.0); n=8 |
| *Dorsal squamodisc* |  |  |  |
| Width | 59.2 ± 11.8 (39.5–89.1); n=23 | 69.1 ± 17.7 (34.2–102.5); n=22 | 99.3 ± 27.7 (55.3–135.4); n=8 |
| Length | 55.5 ± 12.8 (35.9–76.9); n=23 | 57.0 ± 14.6 (26.7–84.2); n=18 | 71.6 ± 14.1 (58.7–103.3); n=8 |
| *Hook* |  |  |  |
| Length | 11.9 ± 1.2 (10.5–16.6); n=24 | 11.0 ± 1.1 (8.4–13.5); n=24 | 13.3 ± 0.6 (12.1–14.0); n=8 |
| *Male copulatory organ (MCO)* |  |  |  |
| Straight length | 52.2 ± 13.3 (32.0–70.7); n=14 | 64.0 ± 10.2 (46.9–88.2); n=27 | 91.5 ± 7.8 (77.2–100.6); n=7 |
| Straight width | 27.4 ± 7.8 (12.4–40.8); n=16 | 36.1 ± 5.6 (19.2–44.9); n=27 | 40.4 ± 3.8 (34.9–45.1); n=7 |
| *Vagina* |  |  |  |
| Total length | 36.3 ± 3.0 (32.9–41.6); n=6 | 43.2 ± 8.2 (32.3–57.8); n=18 | 75.1 ± 18.5 (54.3–97.8); n=4 |
| Tube length | 5.8 ± 1.1 (3.7–7.1); n=7 | 8.5 ± 1.1 (7.2–11.4); n=17 | 17.3 ± 8.3 (7.9–26.8); n=4 |
| Point length | 7.6 ± 2.0 (3.9–10.4); n=8 | 8.3 ± 1.6 (6.4–11.6); n=19 | 11.6 ± 2.0 (10.1–14.5); n=4 |
| *Eyespots* |  |  |  |
| Smaller pair distance | 38.2 ± 9.3 (16.9–49.5); n=28 | 44.8 ± 10.6 (18.6–66.0); n=26 | 95.4 ± 15.6 (67.1–111.6); n=7 |
| Larger pair distance | 20.2 ± 6.7 (6.1–33.1); n=31 | 27.4 ± 10.9 (13.3–65.9); n=28 | 93.3 ± 15.9 (64.8–113.7); n=8 |

Variation in morphology of the measured characteristics was summarised in a biplot through PCA of 17 parameters (Table 2; Fig. 4). The PCA biplot (Fig. 4a) depicts the variation in morphometric measurements where PC1 explains 46.67%, and PC2 explains 14.79% of the variation with main contributors: distance between the eyespots of the larger pair, total body width, total length of the dorsal anchor, maximum width of the ventral bar, and straight length of the ventral bar. The density plot of PC1 scores (Fig. 4b), in accordance with the biplot, indicates a clearly distinctive morphometry in haptoral structures between two morphotypes present in Lake Albert.

**Fig. 4.**
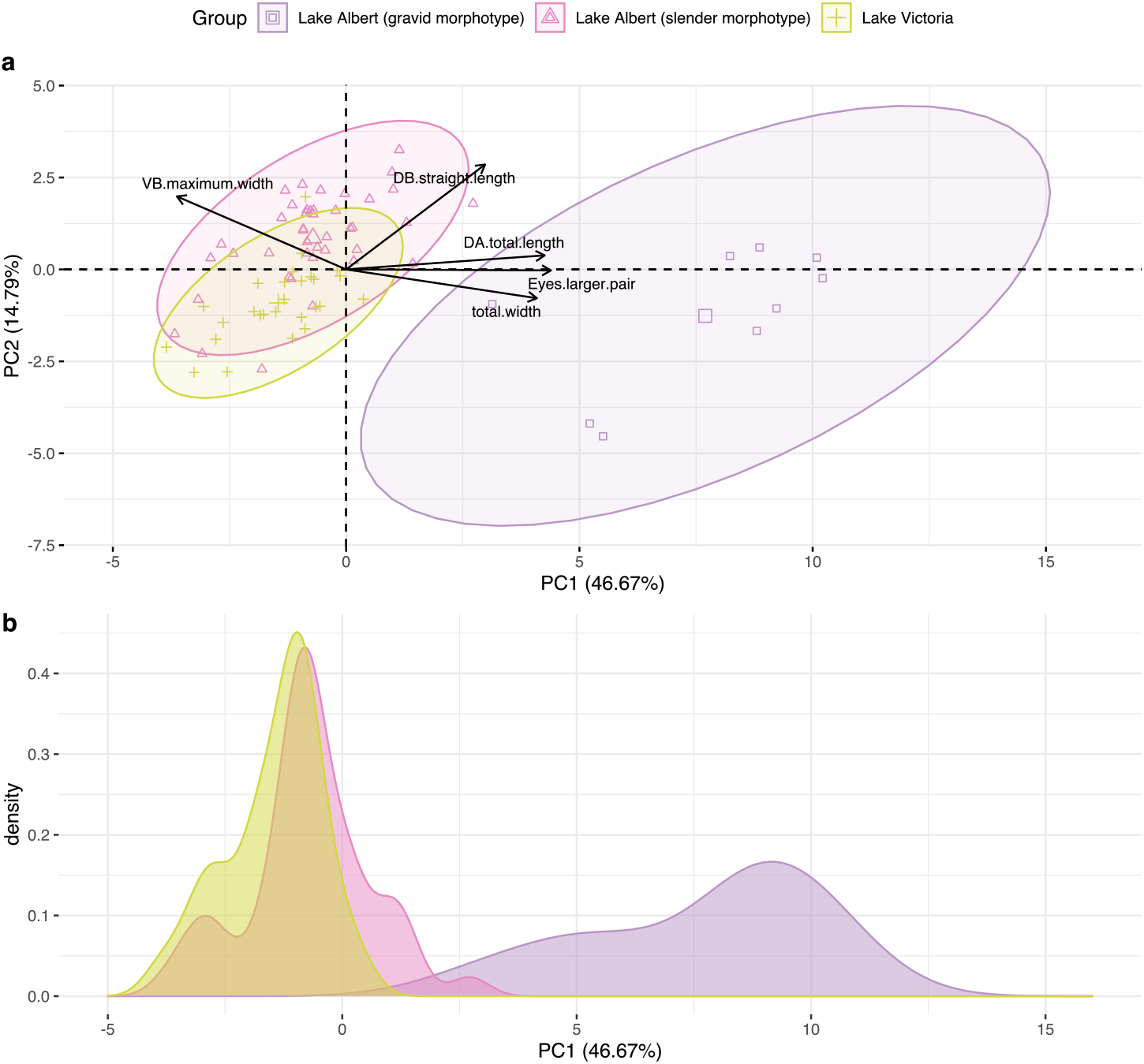
Morphometric variation of haptoral structures and body size in *Dolicirroplectanum lacustre*. (a) Biplot based on PCA from measured characteristics with gravid specimens (n= 9) and slender specimens (n = 42) from Lake Albert, and specimens from Lake Victoria (n= 27) with 95% C.I. ellipses; (b) Density plots depicting the morphometric variation of haptoral structures across populations, summarised in PC1 that explains 46.67% of the haptoral variation. Colours and symbols depict different populations. Vectors indicate the 5 most influential haptoral and body size measurements (DB, dorsal bar; VB, ventral bar; DA, dorsal anchor).

The morphometry of the slender morphotype coincides with the morphometry of the specimens from Lake Victoria (Fig. 4b). We observed similar morphometric variation for the slender morphotype from Lake Albert and the specimens from Lake Victoria. The variation was larger for the gravid morphotype, although this could be an artefact of sample size in Fig. 4 (since the 95 % C.I. is larger when less observations are included for the ellipse). The separation of the gravid morphotype along the PC1 axis from the two other groups can be explained by larger distance between the eyespots of the larger pair, a wider body, and a narrower ventral bar (Fig. 4a, Table 2). MCO straight length (ANOVA, df = 2, *F*= 40.36, p = 0.0000000000028) and straight width (ANOVA, df = 2, *F*= 19.638, p = 0.00000019) were found to differ significantly among the three groups (Fig 5). Overall, MCO length and width were largest for the gravid morphotype (n = 8) from Lake Albert, and smallest in the specimens from Lake Victoria (n = 19). All pairwise comparisons revealed significant differences between the three studied groups (unpaired t-test with Bonferroni correction) (Fig. 5).

**Figure 5.**
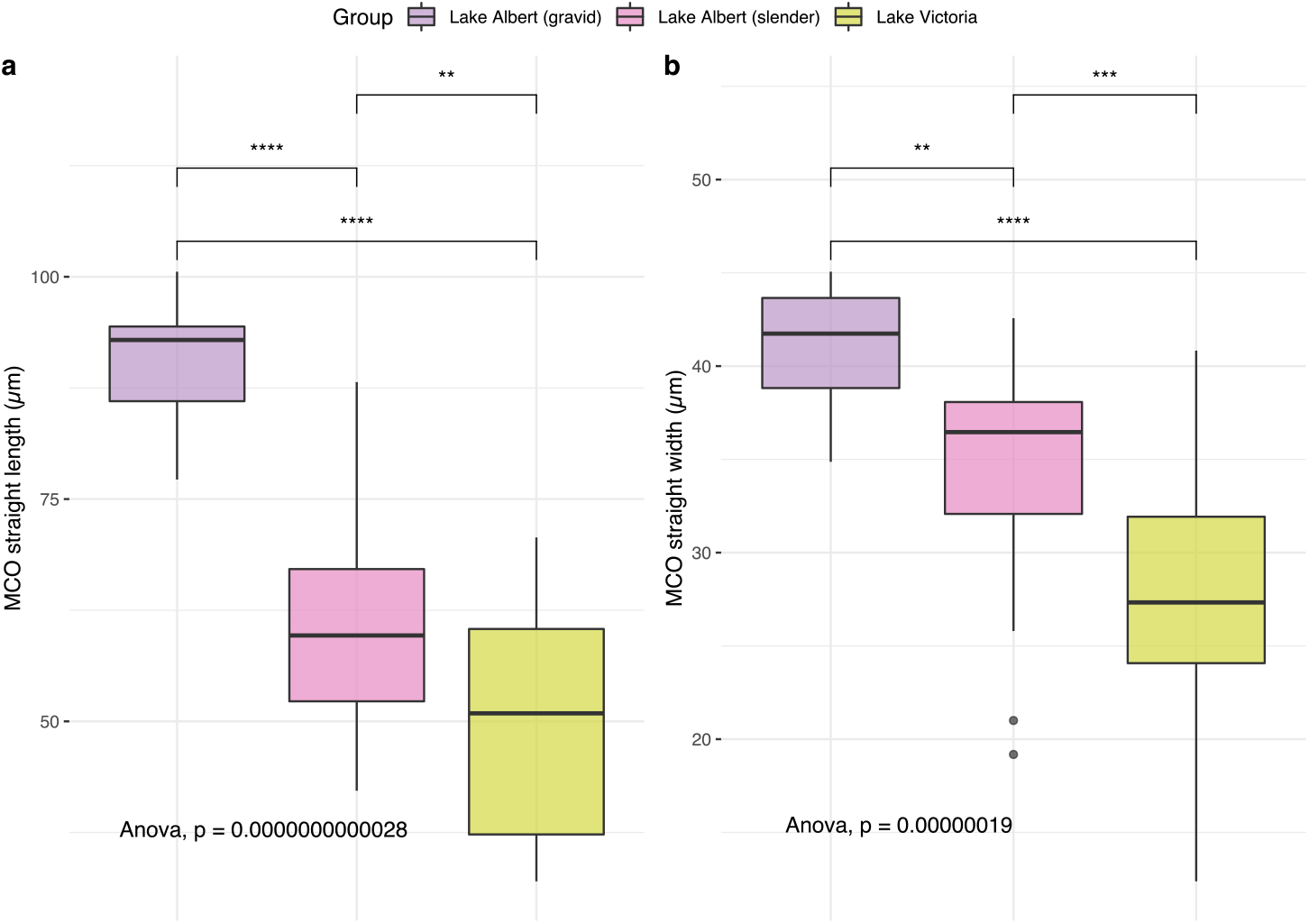
Morphometric variation of the male copulatory organ (MCO) of *Dolicirroplectanum lacustre*. (a) variation in MCO straight length (μm) between morphotypes; (b) variation in MCO straight width (μm) between morphotypes. Colours depict the morphotype from which the MCO measurements were obtained. Following Oneway Anova, the p-values are depicted per MCO measurement, with pairwise comparisons following t-tests with Bonferroni correction for multiple testing (ns, p > 0.05; * p ≤ 0.05; ** p ≤ 0.01; *** p ≤ 0.001; **** p ≤ 0.0001). Box plots with statistical entries: minimum, first quartile, median, third quartile, maximum, outliers. Data from this study combined with measurements of Kmentová et al. (2020a).

### 3.2 Genetic diversity and phylogeography

We found two haplotypes for the 18S rDNA marker (443 bp), two haplotypes for the 28S rDNA marker (778 bp), five haplotypes for the ITS-1 marker (493 bp), and nine haplotypes in the COI marker (325 bp). Uncorrected p-distances varied up to 0.7% for the 18S rDNA marker, and the two haplotypes were shared between lakes and morphotypes. Accordingly, two haplotypes were shared between lakes and morphotypes for 28S rDNA, with uncorrected p-distances up to 2.1%. For the ITS-1 marker, one haplotype was shared between lakes, and uncorrected p-distances varied between 0.2 - 20.6% with a total of 17 indels between haplotypes. Between the morphotypes, no haplotypes were shared for the ITS-1 marker, and uncorrected p-distances varied between 1.5 - 17.2%. For the mitochondrial COI marker, four haplotypes were shared between the lakes, and uncorrected p-distances varied between 0.3 - 13.5%. No haplotypes were shared between the morphotypes, and uncorrected p-distances varied between 12.5 - 13.5% in the COI haplotypes. As all ITS-1 haplotypes could be aligned, our hypothesis that a single diplectanid species was examined is supported (Poisot et al., 2011; Wu et al., 2007). The uncorrected p-distance over the COI fragment does not reach the ‘‘best-compromise threshold” (Meier et al., 2006) for barcoding of 14.5% proposed by Vanhove et al. (2013), which indicates that the specimens are conspecific, and all belong to *D. lacustre*.

We observed no genetic separation between specimens from different lakes. Genetic separation was observed between the morphotypes of *D. lacustre* from Lake Albert in the COI marker, where the gravid morphotype was represented by a single, exclusive haplotype (Fig. 6). There was haplotype sharing between the slender morphotype and the Lake Victoria specimens in the COI marker, and three haplotypes were dominant, surrounded by several satellite haplotypes, separated by single mutations. For the ITS-1 marker, a single haplotype was shared between the gravid population and the Lake Victoria population. The ITS-1 marker did not show the same pattern as the other nuclear markers. Marker incongruence was observed in both lakes. In Lake Albert, one specimen with a gravid morphotype had a COI haplotype dominant in other individuals showing a gravid morphotype, but the nuclear haplotypes were the same as the dominant haplotypes in the slender morphotype. Conversely, in Lake Victoria, two specimens of the slender morphotype and COI haplotype, but with haplotypes of nuclear gene portions corresponding with the gravid morphotype were characterised.

**Fig. 6.**
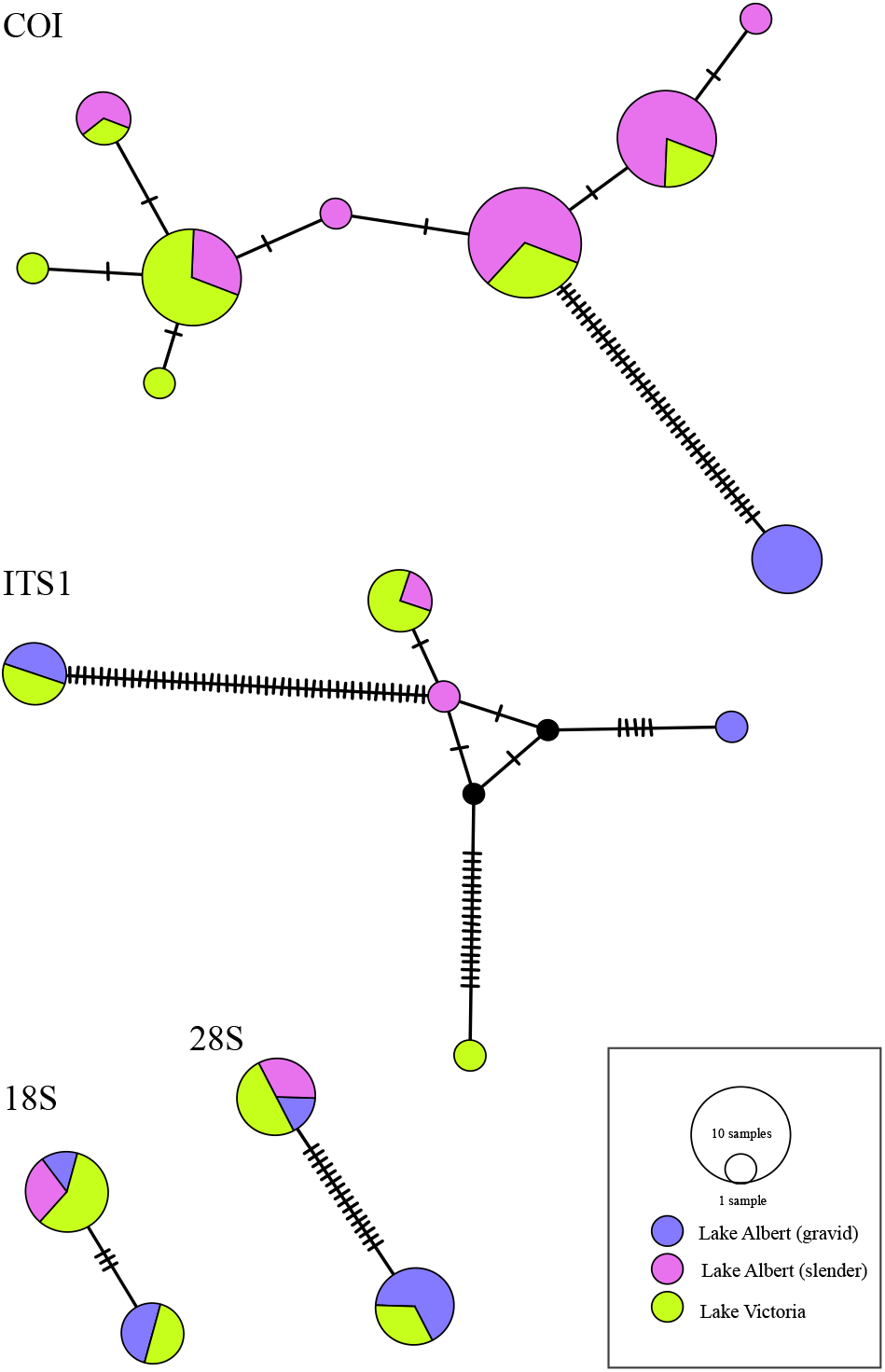
Haplotype networks of *Dolicirroplectanum lacustre* from Lake Albert and Lake Victoria based on sequences of COI mtDNA (n=45), ITS1 rDNA (n=11), 28S rDNA (n=12), 18S rDNA (n=11) combining data generated during this study and by Kmentová et al. (2020a). Circles represent different haplotypes with the size proportional to the number of individuals sharing the haplotype. Colours correspond to the different populations. Mutation steps as hatch marks.

The genetic diversity in the COI mtDNA gene portion was largest in Lake Albert (Table 3). The number of polymorphic sites in COI was 46 (n=29) in Lake Albert, and six (n=16) in Lake Victoria. Five sites were polymorphic (n=24) in the slender morphotype, and for the gravid morphotype only a single haplotype was represented in five individuals. We observed similar levels of nucleotide and haplotype diversity between *D. lacustre* in both lakes, however values for these diversity measures were larger in Lake Albert. F_ST_ values were significant between the Lake Albert morphotypes (F_ST_= 0.97049, p-value = 0.00000). Likewise, the gravid morphotype and the specimens from Lake Victoria were genetically differentiated (F_ST_= 0.97024, p-value = 0.00000). Conversely, low F_ST_ values (F_ST_= 0.16794, p-value = 0.01802) for the Lake Victoria specimens and the slender specimens from Lake Albert indicate higher genetic similarity between these populations. Following AMOVA between specimens from Lake Albert and Lake Victoria (F_ST_= 0.05983, p-value = 0.03812), most of the variation was present within the lakes (94.02%) in comparison with 5.98% among-lake variation.

**Table 3.** Genetic diversity indices of *Dolicirroplectanum lacustre* in Lake Albert (per population) and Lake Victoria inferred from a 325 bp portion of the mitochondrial cytochrome *c* oxidase subunit I (COI) region.

| Population | *n* | H | S | Maximum uncorrected p-distance | Hd | $\pi$ |
| --- | --- | --- | --- | --- | --- | --- |
| Lake Albert | 29 | 7 | 46 | 13.2 % | 0.8079 ± 0.0405 | 0.041228 ± 0.021246 |
| Lake Victoria | 16 | 6 | 6 | 1.2 % | 0.7667 ± 0.0839 | 0.004949 ± 0.003462 |
| Lake Albert (slender) | 24 | 6 | 5 | 1.5 % | 0.7536 ± 0.0562 | 0.004515 ± 0.003159 |
| Lake Albert (gravid) | 5 | 1 | 0 | / | / | / |

*n*, sample size; H, number of haplotypes; S, number of polymorphic sites; Hd, haplotype diversity; $\pi$ , nucleotide diversity

### 3.3 Demographic history

No signatures of population expansion could be detected for *D. lacustre* from the studied lakes. Neutrality test statistics were not significant for Lake Albert alone (Tajima’s D = 0.537, p = 0.774; Fu’s F_S_= 9.872, p = 0.995), as for Lake Albert and Lake Victoria combined (Tajima’s D = −0.230, p = 0.453; Fu’s F_S_= 7.137, p =0.981).

### 3.4 The gill microhabitats

Individuals of *D. lacustre* were predominantly attached to the median-central and median-distal microhabitats on the gills (Fig. 7). In Lake Victoria, parasites were mostly retrieved from the right gill chamber (2 left, 18 right). In Lake Albert, most specimens were retrieved from the left gill chamber (slender: 50 left, 22 right; gravid: 8 left). In co-infections, both morphotypes occupied the same microhabitats, but several specimens of the gravid morphotype were mostly found on the median-ventral portion of the gill (Fig. 7). Whereas specimens from Lake Victoria were attached to the proximal-ventral portion of the gill, this was uncommon in Lake Albert.

**Fig. 7.**
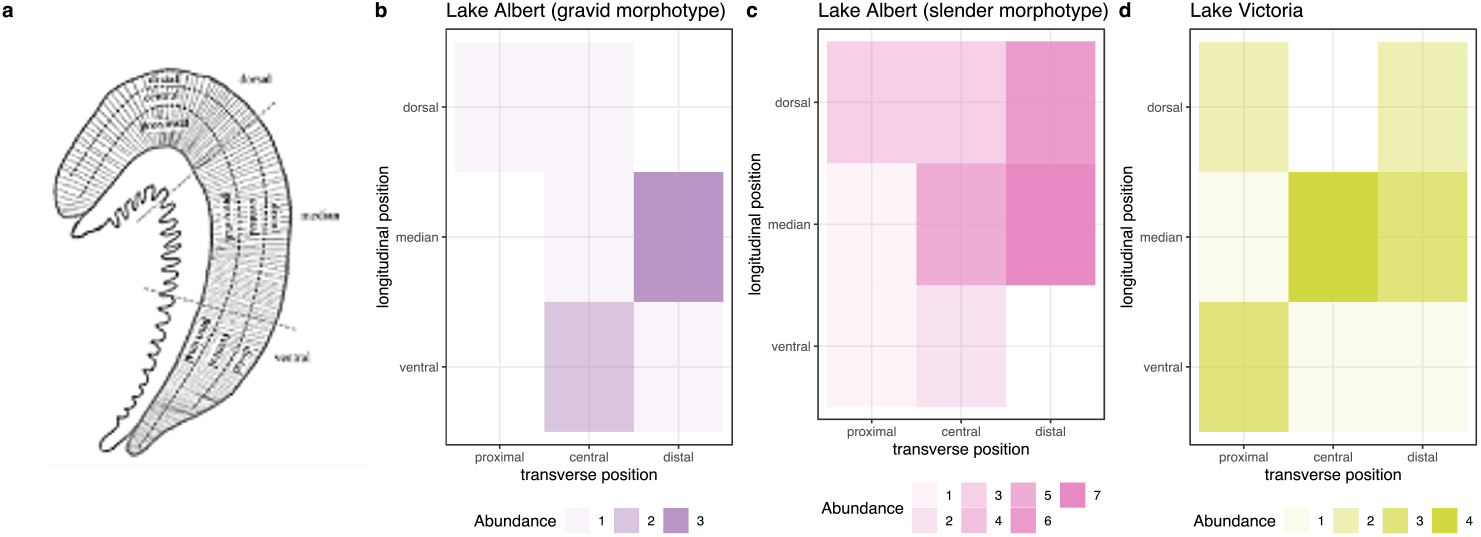
Microhabitat distribution of *Dolicirroplectanum lacustre* visualised in heat plots. (a) division of the fish gill into nine microhabitats across a longitudinal and transversal axis; (b) Lake Victoria; (c) gravid morphotype from Lake Albert; (d) slender morphotype from Lake Albert.

## 4. Discussion

In Lake Albert we morphologically identified two distinct morphotypes of *Dolicirroplectanum lacustre*, a monogenean parasite infecting latid fishes across the African freshwater system. A single morphotype (slender) is suggested to be co-introduced in Lake Victoria with the historical introductions of the Nile perch. The variation in haptoral structures of the slender morphotype coincides with the variation in specimens from Lake Victoria, with the latter showing slightly less phenotypic and genetic variation. We observed spatial similarity in the microhabitat distribution between the specimens of the slender morphotype from both lakes. Co-infections with both morphotypes were observed in Lake Albert. The two morphotypes were genetically separated in the COI mtDNA region, but genetic differentiation did not meet the level typically associated with species-delineation. Patterns in the ITS-1 region differed in the slender specimens between Lake Victoria and Lake Albert. Mitonuclear marker incongruence was observed within the morphotypes of *D. lacustre*.

### A host species harbours the greatest diversity of parasites in the area where it resided longest

The haplotype sharing between studied locations supports the co-introduction of *D. lacustre* with *L. niloticus* to Lake Victoria from Lake Albert (Pringle, 2005). This agrees with host data provided by Hauser et al. (1998), which confirmed the presence of *L. niloticus* from Lake Albert in Lake Victoria and excluded the successful establishment of introduced Nile perch from Lake Turkana. We observed large variation in haptoral morphology in Lake Albert (native range) and identified the two morphotypes as in earlier studies (Kmentová et al., 2020a; Thurston and Paperna, 1969). The presence of different species of Nile perch (Hauser et al., 1998), that occur in sympatry in Lake Albert may have provided *D*. *lacustre* with different gill habitats and could account for the morphological variation between morphotypes in Lake Albert. Under this scenario, the observations of co-infections would have required secondary contact and host switching or hybridisation between the two host taxa. The reduced phenotypic variation (and presence of a single morphotype) in Lake Victoria may be an indication for a founder effect. In accordance, we have observed reduced genetic diversity in the specimens from Lake Victoria in comparison with Lake Albert. When relatively few parasite specimens were co-introduced, a ‘founder effect’ would be expected (Mayr, 1942). Accordingly, a host species is expected to harbour the greatest diversity of parasites in the area where it resided longest (Manter 1966). This argument is supported by the low number of introduced hosts (n = 382) into Lake Victoria (Hauser et al., 1998).

### Phenotypic plasticity: discordant variation and mitonuclear discordance

The observed variation in the morphology and resulting morphotypes, in combination with below species-level genetic differentiation, suggests phenotypic plasticity of the species *D. lacustre* following the definition of DeWitt et al. (1998): species with “the potential to produce a range of different, relatively fit phenotypes in multiple environments”. Discordance between molecular and morphological differentiation has been observed for other diplectanid monogenean species, namely *Lamellodiscus* spp. (Diplectanidae) (Desdevises et al., 2000; Poisot et al., 2011). Following Desdevises et al. (2000), these inconsistencies could be attributed to phenotypic variability of some characters related to the environment and the host (Villar-Torres et al. 2019). Moreover, the observed phenotypic variability and the genetic structuring of the morphotypes observed in the COI marker could be an indication for ongoing sympatric speciation or diversification processes (Poulin, 2002) of *D. lacustre* in Lake Albert. Parasites with direct life cycles, like monogeneans, are expected to experience more favourable conditions for sympatric speciation, especially when infecting long-living hosts (Brooks and McLennan, 1993), such as the Nile perch. Similarly, the observed morphological differentiation in the MCO could be an indication of ongoing diversification or incipient speciation (Šimková et al., 2002), explained by the ‘reinforcement hypothesis’ (Rohde, 1991): “reinforcement of reproductive barriers is one of the main factors resulting in niche segregation”.

Although the presented population structure based on the COI marker indeed agrees with the reproductive isolation of the gravid morphotype, the mitonuclear discordance gives a different picture, and indicates an incomplete reproductive barrier between the two morphotypes. Potentially, this discordance may be explained by intraspecific gene flow by hybridisation between the morphotypes in Lake Albert, or incomplete lineage sorting (Després 2019). The former has earlier been linked with the evolution of invasiveness (Culley and Hardiman, 2008).

### A ‘failure to diverge’ in Lake Tanganyika versus ongoing diversification in Lake Albert

Although former studies identified two different morphotypes of *D. lacustre* in Lake Albert, the fine-scale genetic differentiation between the morphotypes had not been investigated. In Kmentová et al. (2020a), a ‘failure to diverge’ was observed for *D. lacustre*, as the species had not diversified over the variety of African freshwater systems on a wide geographic scale, even when infecting different host species in Lake Tanganyika. This was hypothesised to be attributed to a low rate of molecular evolution. Our data supports the severely reduced genetic diversity in Lake Tanganyika in comparison to the other studied areas (Supplementary table S1). Moreover, the haplotype network for COI mtDNA indicates that the gravid lineage from Lake Albert is genetically more distinctive from the slender lineage in Lake Albert and Lake Victoria, compared to the specimens from Lake Tanganyika (Supplementary figure S2). The reduced diversification in *D. lacustre* from Lake Tanganyika may be explained by the recently estimated timing of colonisation by *L. niloticus* and divergence (1.27-1.76 MYA) of the four extant *Lates* spp. in Lake Tanganyika (Koblmüller et al., 2021).

### Future perspectives

Although this study sheds light on the population-scale differentiation and the co-introduction of *D. lacustre*, several questions on its co-introduction and diversification mechanisms remain. We suggest: (1) to investigate whether the diversification patterns after co-introduction observed in this study, occur across different freshwater systems, we should increase the number of samples over various sampling sites within the entire native range (including Lake Turkana) and introduced range (Lake Kyoga) of Nile perch. By sample acquisition from historical collections, broader spatial and temporal patterns of diversification after co-introduction could be examined; (2) with sample collection note should be taken of the host (species) identity, in particular in Lakes Albert and Turkana, where two sympatric species could occur, since host identity may contribute to the observed diversification patterns of *D. lacustre;* (3) the application of Next Generation Sequencing approaches to study genome-wide intraspecific variation in *D. lacustre*, in order to identify the determinants of the observed diversity and mitonuclear discordance; (4) to undertake a more detailed study on the gill environment of Nile perch and microhabitat selection of *D. lacustre* in Lake Albert to determine whether microhabitat preference plays a role in the ongoing morphological and genetic diversification and possible sympatric speciation of *D. lacustre*.

## Acknowledgments

The authors would like to thank the Zoology Research group: Biodiversity & Toxicology at Hasselt University. Natascha Steffanie is acknowledged for sample processing, and Armando Cruz Laufer is thanked for laboratory assistance. We acknowledge the Royal Belgian Institute of Natural Sciences and the Royal Museum for Central Africa for sample provision.

## Statements & Declarations

### Funding

Study supported by Special Research Fund (BOF) UHasselt: BOF21DOC08 (KJMT), BOF20TT06 (MPMV), and BOF21PD01 (NK); by Czech Science Foundation standard project GA19-13573S; and by Research Foundation - Flanders (FWO-Vlaanderen) research grant 1513419N; infrastructure funded by EMBRC Belgium - FWO project GOH3817N. The funders had no role in study design, data collection and analysis, decision to publish, or preparation of the manuscript. We thank the Royal Belgian Institute of Natural Sciences and Royal Museum for Central Africa for sample collection under the BELSPO Brain project, HIPE (BR/154/A1/HIPE).

### Author contributions

All authors contributed to the study conception and design. Material preparation, and data collection were performed by Kelly J.M. Thys, Jonas W.J. Custers, Nikol Kmentová, Nathan Vranken, and Maarten Van Steenberge. Kelly J.M. Thys and Jonas W.J. Custers conducted data analysis. The original draft of the manuscript was written by Kelly J.M. Thys, and revision of the previous versions of the manuscript was provided by Nikol Kmentová, Maarten Van Steenberge, and Maarten P.M. Vanhove. Funding was acquired by Kelly J.M. Thys, Nikol Kmentová, Maarten Van Steenberge, and Maarten P.M. Vanhove. All authors read and approved the final manuscript.

### Compliance with Ethical Standards

The authors declare that they have no competing interests.

### Data availability

All data generated or analysed during this study are included in this published article [and its supplementary information files].

## Supplementary data

**Supplementary table S1.** Genetic diversity indices of *Dolicirroplectanum lacustre* in Lake Albert (per population) and Lake Victoria inferred from a 325 bp portion of the mitochondrial cytochrome c oxidase subunit I (COI) region.

| Population | *n* | H | S | Maximum uncorrected p-distance | Hd | $\pi$ |
| --- | --- | --- | --- | --- | --- | --- |
| Lake Albert | 29 | 7 | 46 | 13.2 % | 0.8079 ± 0.0405 | 0.041228 ± 0.021246 |
| Lake Victoria | 16 | 6 | 6 | 1.2 % | 0.7667 ± 0.0839 | 0.004949 ± 0.003462 |
| Lake Albert slender | 24 | 6 | 5 | 1.5 % | 0.7536 ± 0.0562 | 0.004515 ± 0.003159 |
| Lake Albert gravid | 5 | 1 | 0 | / | / | / |
| Lake Tanganyika | 38 | 3 | 2 | 0.3 % | 0.2404 ± 0.0858 | 0.000757 ± 0.000976 |

*n*, sample size; H, number of haplotypes; S, number of polymorphic sites; Hd, haplotype diversity; $\pi$ , nucleotide diversity

**Supplementary figure S2.**
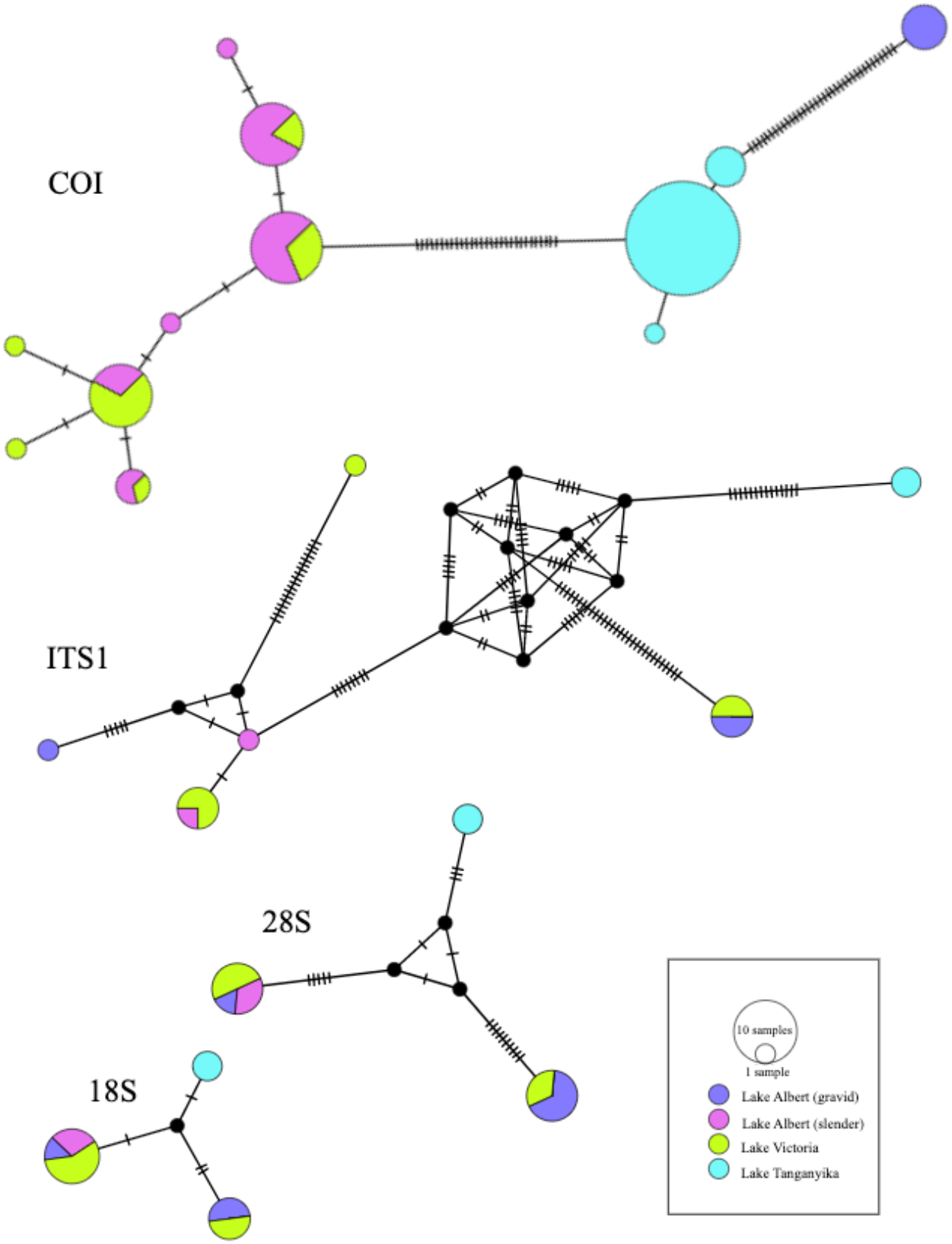
Haplotype networks of *Dolicirroplectanum lacustre* from Lake Albert, Lake Victoria and Lake Tanganyika based on COI mtDNA sequences (n=83), ITS1 rDNA (n=13), 28S rDNA (n=14), 18S rDNA (n=13) combining data generated during this study and Kmentová et al. (2020a). Circles represent different haplotypes with the size proportional to the number of individuals sharing the haplotype. Colours correspond to the different populations. Mutation steps as hatch marks.

## Notes

### Competing Interest Statement

The authors have declared no competing interest.

### Summary of Updates

Submission to International Journal for Parasitology

